## Supplementary figures and legends for "Capsid structure of a metazoan fungal dsRNA megabirnavirus reveals its uniquely acquired structures"

### Supplementary figures and figure legends

| Parameter or statistic | Full RnMBV1 | Asymmetric full RnMBV1 | Empty RnMBV1 |
|---|---|---|---|
| Data collection and processing | | | |
| Magnification | 59,000 | 59,000 | 59,000 |
| Voltage (kV) | 300 | 300 | 300 |
| Electron exposure ($e^-/Å^2$) | 48 | 48 | 48 |
| Defocus range (μm) | -1.00 to -2.75 | -1.00 to -2.75 | -1.00 to -2.75 |
| Pixel size (Å) | 1.12 | 1.12 | 1.12 |
| Symmetry imposed | I | C1 | I |
| Initial micrographs (No.) | 2,839 | 2,839 | 2,839 |
| Final micrographs (No.) | 2,734 | 2,734 | 2,734 |
| Initial particle images (No.) | 45,869 | 45,869 | 45,869 |
| Final particle images (No.) | 12,230 | 244,609 | 14,602 |
| Map resolution (Å) | 3.2 | 3.3 | 3.2 |
| FSC threshold | 0.143 | 0.143 | 0.143 |
| Map resolution range (Å) | 3.1-4.1 | 3.1-5.2 | - |
| Refinement | | | |
| Map sharpening *B* factor ($Å^2$) | -173 | -173 | -179 |
| Model composition | | | |
| Non-hydrogen atoms | 18,680 | 996 | - |
| Protein residues | 2,418 | 135 | - |
| *B* factors ($Å^2$) | | | |
| Protein (min./max./mean) | 9.7/68.59/23.81 | 35.38/52.98/41.81 | -/-/- |
| RMSD* | | | |
| Bond lengths (Å) | 0.007 | 0.005 | - |
| Bond angles (°) | 0.649 | 1.051 | - |
| Validation | | | |
| MolProbity score | 1.82 | 1.98 | - |
| Clashscore | 8.12 | 6.98 | - |
| Poor rotamers (%) | 0 | 0 | - |
| Ramachandran plot | | | |
| Favored (%) | 94.49 | 87.97 | - |
| Allowed (%) | 5.51 | 12.03 | - |
| Disallowed (%) | 0 | 0 | - |
| CC (mask) | 0.89 | -0.03 | - |
| EMRinger score | 4.02 | 2.04 | - |
| Data deposition | | | |
| EMDB | EMD-15855 | EMD-15859 | EMD-15857 |
| PDB | 8B4Z | 8B59 | - |

*RMSD, root mean square deviation

**Supplementary Table. S1 Cryo-EM data collection, refinement, and validation statistics.**

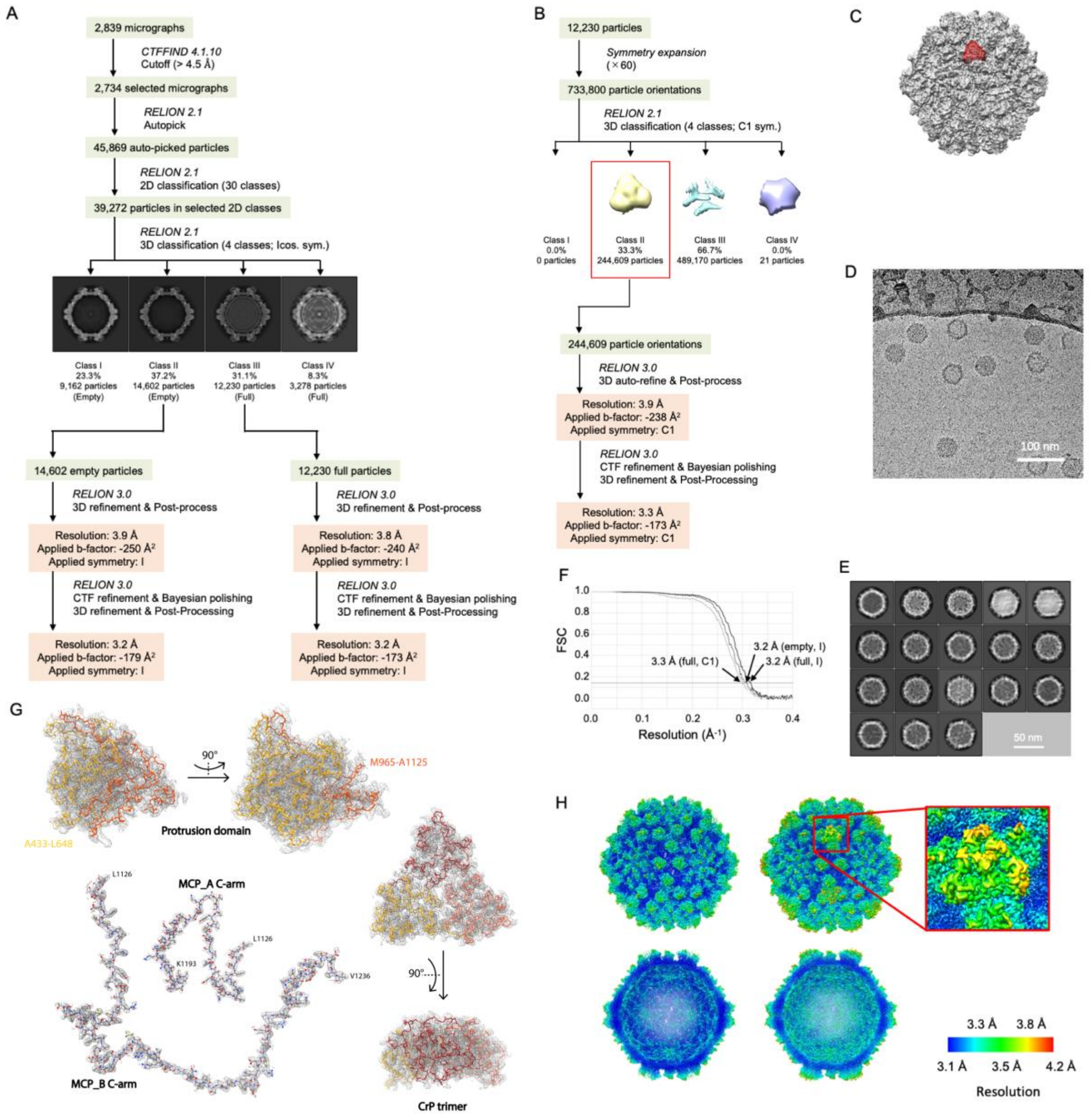


**Supplementary Fig. S1 Summary of the cryo-EM data analysis.** A) Flowchart of the RnMBV1 MCP data analyses of the cryo-EM micrographs using RELION. B) Flowchart on generating the RnMBV1 CrP 3D reconstruction using RELION. C) The CrP's location on the RnMBV1 capsid on the map. D) The cryo-EM raw image of the RnMBV1 particles (empty/full). E) The 2D class overview. F) The gold-standard FSC resolution curves of the full particle in icosahedral (I) symmetry (solid black line), empty particle in symmetry I (dashed line), and full particle in C1 symmetry (gray line) reconstructions for RnMBV1 virions. The resolutions of these cryo-EM models are estimated as 3.2,

3.2, and 3.3 Å, respectively (FSC cutoff = 0.143). G) The cryo-EM map and fitting with the atomic models of the protrusion domain (yellow and orange shown in backbone representation only), the C-terminal arms of the MCPs (in ball-and-stick mode), and the CrPs trimer (red, orange, and dark yellow shown in backbone representation). H) Overall 3D reconstructions. Top panel: The overall capsid map with or without the presence of CrP reconstructions. Bottom panel: The back half of the capsid 3D reconstructions shows the interior features with or without the presence of CrP occupancy. The capsid is colored from blue to red, in accordance with the estimated local resolutions.

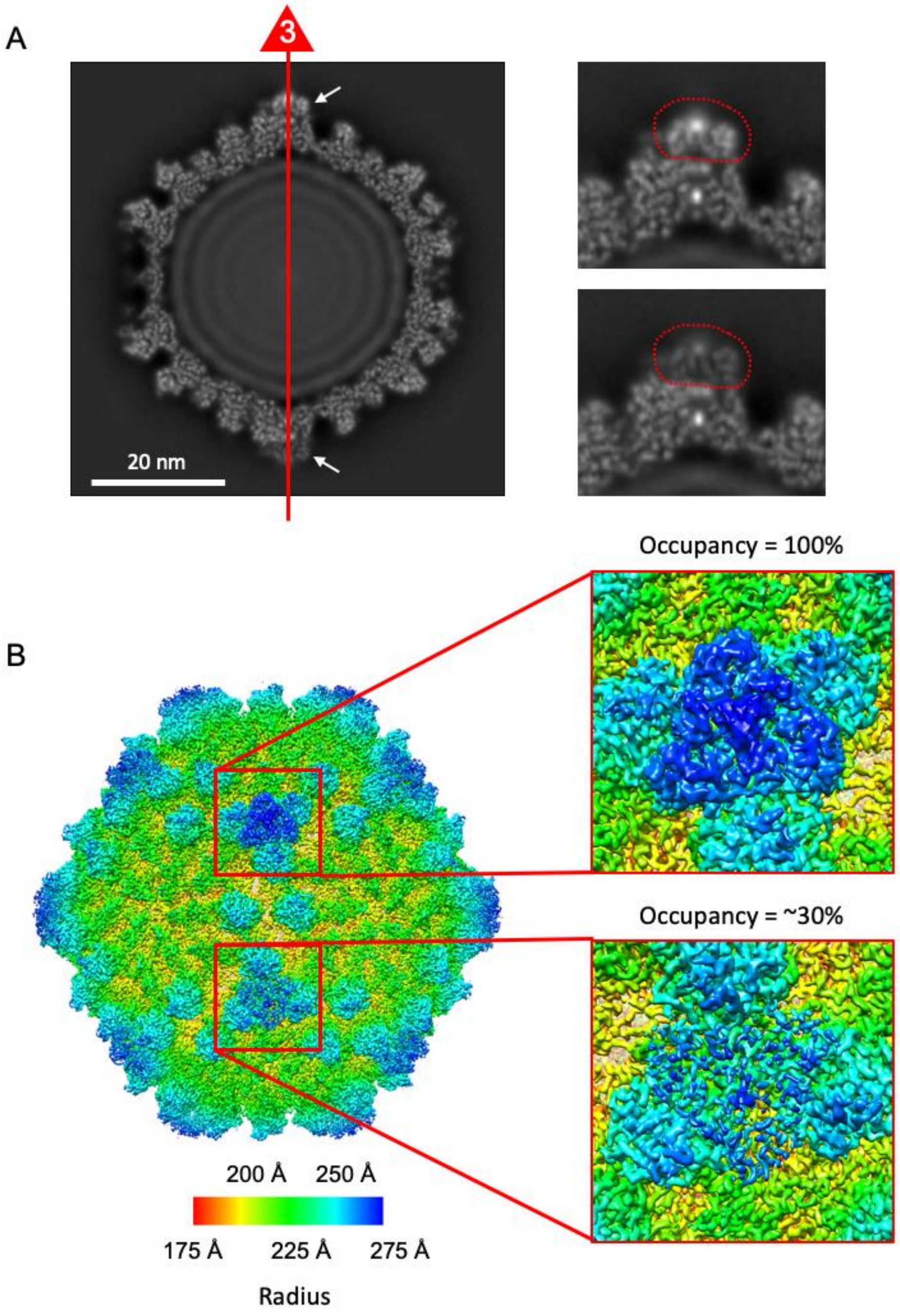


**Supplementary Fig. S2 Reconstructed cryo-EM map after the 3D focused classification on CrP trimers.** A) Overall central cross-section of the RnMBV1 cryo-EM map viewed along the icosahedral 3-fold axis (left panel). Scale bar: 20 nm. Enlarged views of the CrPs (right panels). The CrP trimer is surrounded by a red dotted line. The map intensity level in one CrP trimer (lower right panel) is weaker than that in the other regions, while the map intensity in another CrP trimer (upper right panel) is as strong as that in the other regions. B) Surface representation of the reconstructed cryo-EM map. The intensity level in one CrP trimer is recovered as high after the 3D focused classification, which is deemed to be 100% occupancy (right upper panel). The map intensity in the other CrP trimers is low, which corresponds to approximately 30% occupancy (right bottom panel). The image is colored radially according to the distance from the particle center.

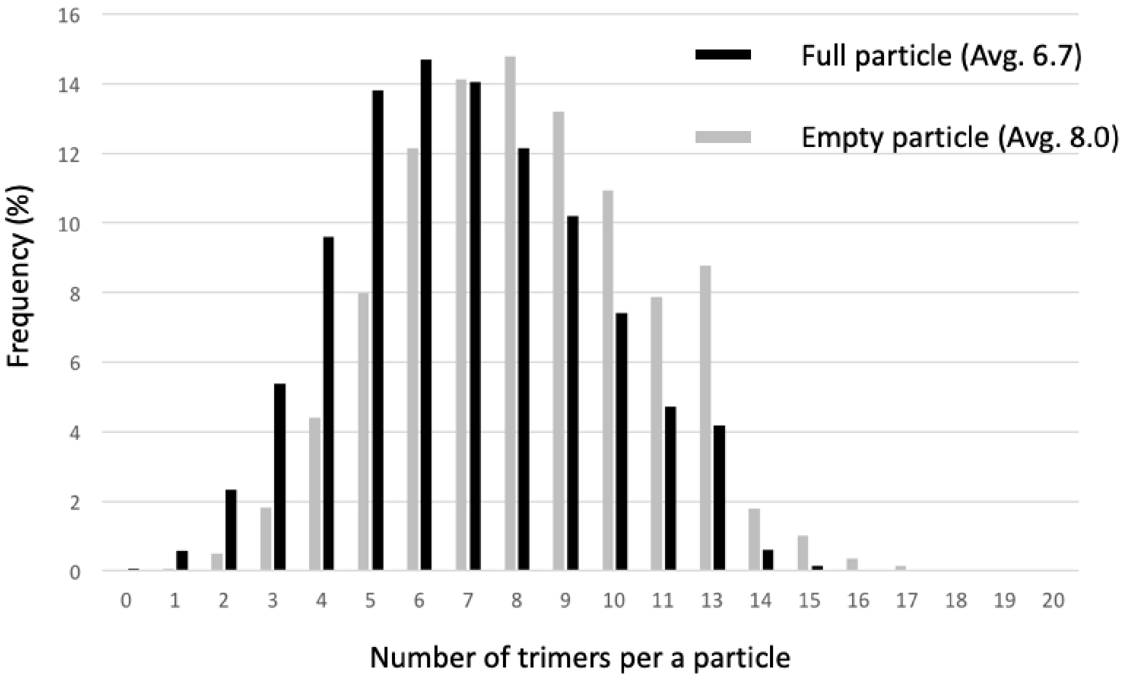


**Supplementary Fig. S3 Number of CrP trimers on the particle and the distribution.** After the 3D focused classification of CrP trimers, the number of CrP trimers bound to the particle is counted and the distribution is plotted. The averaged numbers of presented CrP trimers per particle are 6.7 and 8.0 for the full and empty particles, respectively.

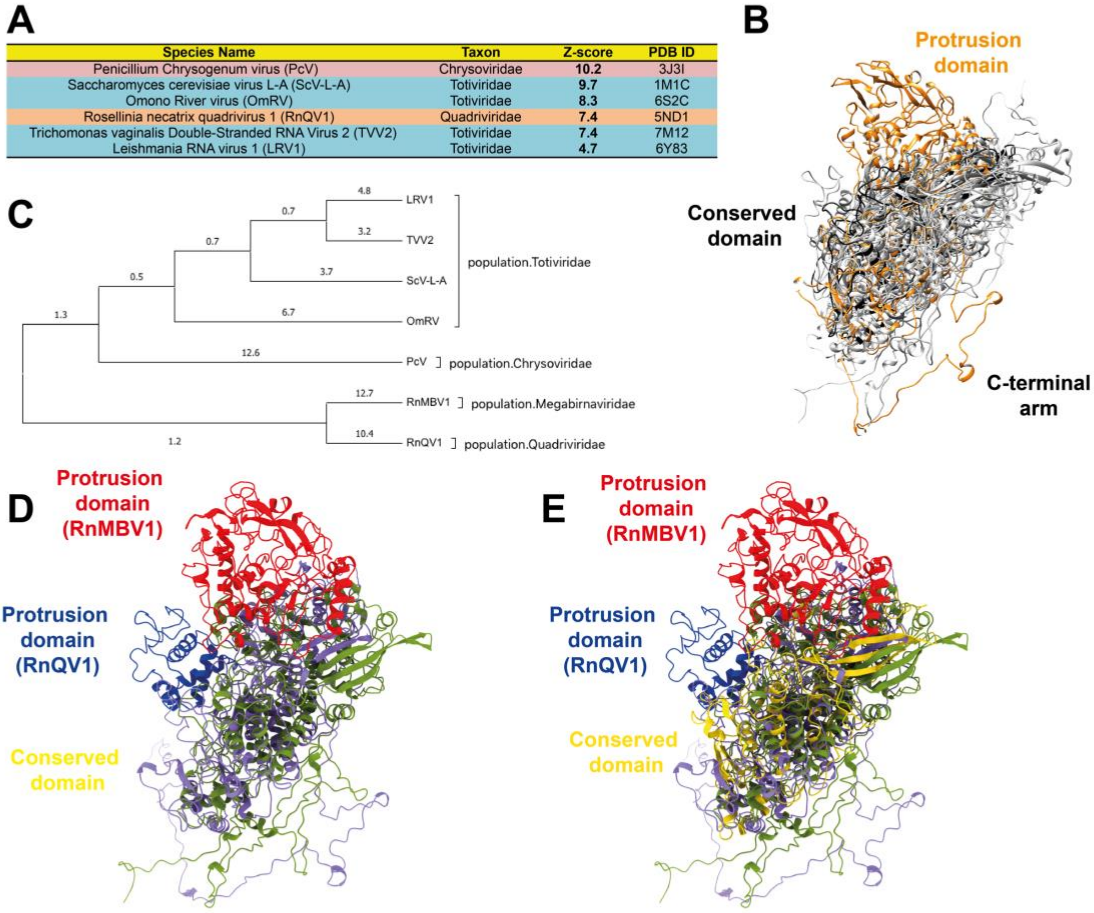


| Species Name | Taxon | Z-score | PDB ID |
|---|---|---|---|
| Penicillium Chrysogenum virus (PcV) | Chrysoviridae | 10.2 | 3J3I |
| Saccharomyces cerevisiae virus L-A (ScV-L-A) | Totiviridae | 9.7 | 1M1C |
| Omono River virus (OmRV) | Totiviridae | 8.3 | 6S2C |
| Rosellinia necatrix quadrivirus 1 (RnQV1) | Quadriviridae | 7.4 | 5ND1 |
| Trichomonas vaginalis Double-Stranded RNA Virus 2 (TVV2) | Totiviridae | 7.4 | 7M12 |
| Leishmania RNA virus 1 (LRV1) | Totiviridae | 4.7 | 6Y83 |

**Supplementary Fig. S4 Structural alignments, phylogeny, and superimpositions of the RnMBV1 MCP and the related CP of dsRNA viruses.** A) The summary table of the viral CP structures similar to that of the RnMBV1 MCP identified using the Dali search. The background colors indicate different taxonomy clades. B) Superimpositions of the RnMBV1 MCP (orange) and the six RnMBV1-related CPs listed in A) (gray). C) Structure-based phylogenetic tree of the RnMBV1 MCP and the structurally close-related six CPs. D) The superimposition of the RnMBV1 MCP (purple and red) and RnQV1 CP (green and dark blue). E) The superimposition of RnMBV1 MCP (purple and red), RnQV1 CP (green and dark blue), and yeast ScV-L-A CP (yellow). D, E) The red and blue portions show the protrusion domain in the MCP of RnMBV1 and the CP of RnQV1, respectively.

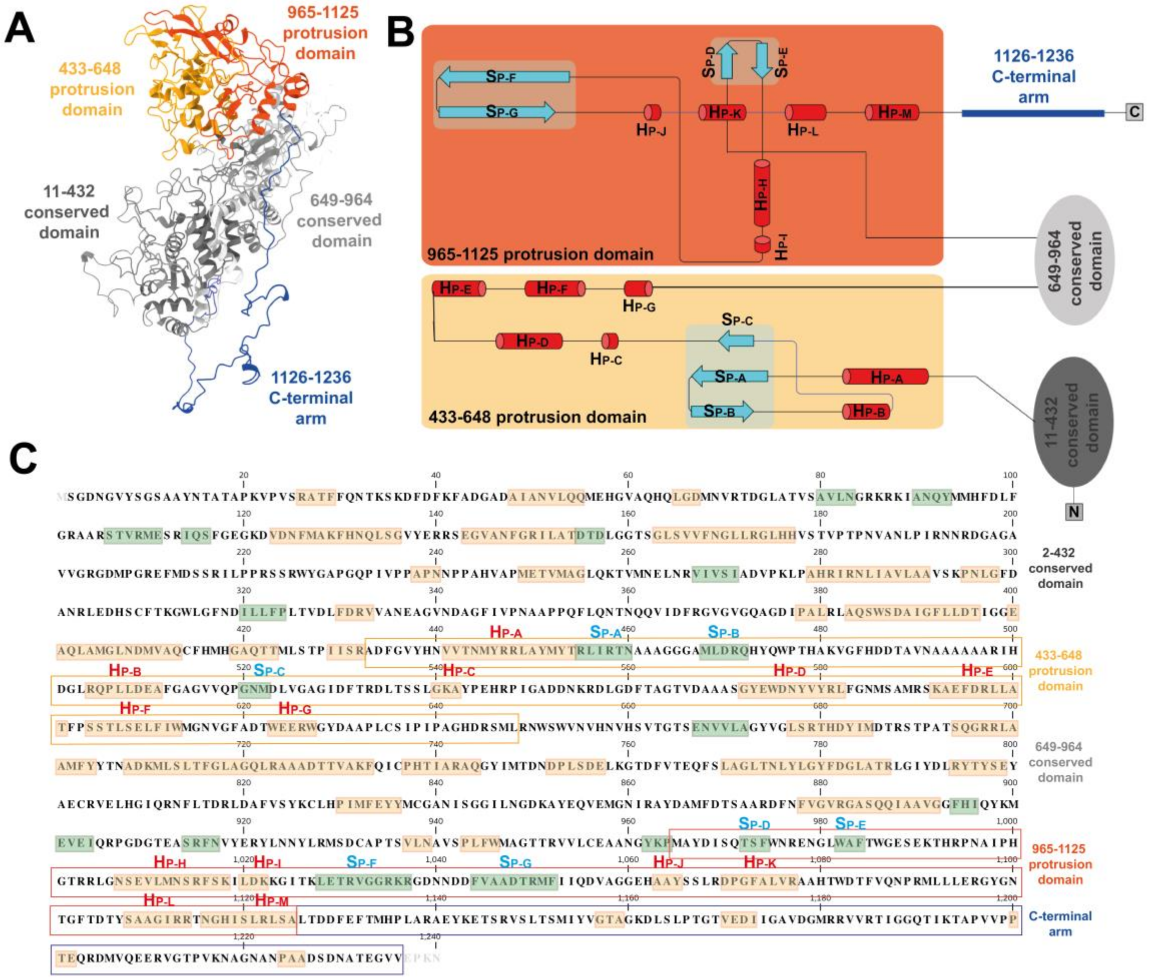


**Supplementary Fig. S5 Atomic model, structural topology, and secondary structural elements of MCP.** A) The atomic model of one MCP colored by domains. The protrusion domains (amino acid residues 433–648 and 965–1125) are colored yellow and orange, respectively and the conserved domains (amino acid residues 11–432 and 649–964) are colored dark gray and light gray. The C-terminal arm (amino acid residues 1126–1236) is colored dark blue. B) Structural topology diagram of the MCP. The color codes correspond to those in A). The α-helices and β-strands are shown as red cylinders and light blue arrows and named accordingly. The blue, translucent, rounded boxes cover the regions containing β-sheets. C) The amino acid sequence organization of the MCP. The orange and green boxes highlight α-helices and β-strands, respectively. The colors and labels correspond to those of B).

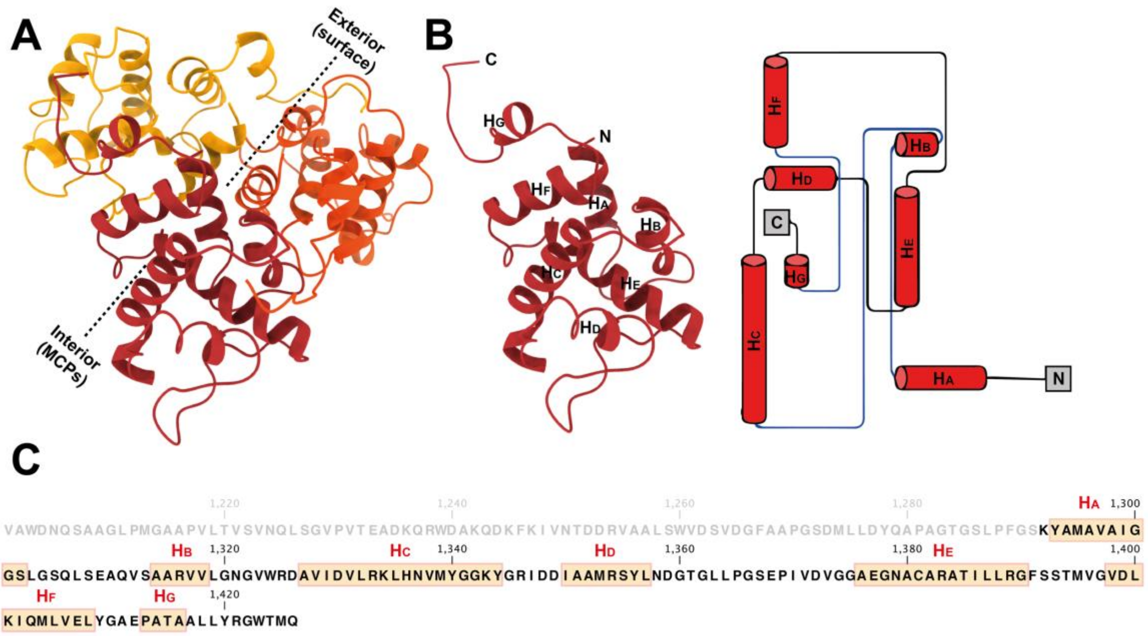


**Supplementary Fig. S6 Atomic model, structural topology, and secondary structural elements of CrP.** A) The atomic model of one CrP trimer (red, orange, and dark yellow). The dashed lines indicate the exterior and interior sides of the CrP trimer. B) Structural topology diagram of the CrP. All α-helices are colored red and numbered in order (assigned to be $H_{A\text{-}G}$ from the N- to C-terminus). C) The amino acid sequence organization of CrP. The amino acid sequence is a C-terminal portion of RnMBV1 ORF3. The amino acid residues of CrP are marked in black bold from Lys1292 to Gln1426. The orange boxes highlight the α-helices.
