## Supplementary material for "Capsid structure of a metazoan fungal dsRNA megabirnavirus reveals its uniquely acquired structures": Funding information

Funding was provided by the following agencies: Vetenskapsrådet (VR)/The Swedish Research Council (to K.O., grant no. 2018-03387), FORMAS research grant from the Swedish Research Council for Environment, Agricultural Sciences, and Spatial Planning (to K.O., grant no. 2018-00421), and the Royal Swedish Academy of Sciences (to K.O., grant no. BS2018-0053). Grants-in-Aid for Scientific Research on Innovative Areas from the Japanese Ministry of Education, Culture, Sports, Science and Technology (KAKENHI 21H05035, 17H01463, 16H06436, 16H06429 and 16K21723 to N.S., 15K18521 and 18K06154 to N.M.), the Collaborative Study Program of the National Institute for Physiological Science (to N.M.), the Platform Project for Drug Discovery, Informatics, and Structural Life Science (PDIS) from the Ministry of Education, Culture, Sports, Science and Technology (MEXT) and for Supporting Drug Discovery and Life Science Research (Basis for Supporting Innovative Drug Discovery and Life Science Research (BINDS)) (to N.M.).
